# ScaleQC: A Scalable Lossy to Lossless Solution for NGS Sequencing Data Compression

**DOI:** 10.1101/2020.02.09.940932

**Authors:** Rogshan Yu, Wenxian Yang

## Abstract

**Motivation:** Per-base quality values in NGS sequencing data take a significant portion of storage even after compression. Lossy compression technologies could further reduce the space used by quality values. However, in many applications lossless compression is still desired. Hence, sequencing data in multiple file formats have to be prepared for different applications.

**Results:** We developed a scalable lossy to lossless compression solution for quality values named ScaleQC. ScaleQC is able to provide bit-stream level scalability. More specifically, the losslessly compressed bit-stream by ScaleQC can be further truncated to lower data rates without re-encoding. Despite its scalability, ScaleQC still achieves same or better compression performance at both lossless and lossy data rates compared to the state-of-the-art lossless or lossy compressors.

**Availability:** ScaleQC has been integrated with SAMtools as a special quality value encoding mode for CRAM. Its source codes can be obtained from our integrated SAMtools (https://github.com/xmuyulab/samtools) with dependency on integrated HTSlib (https://github.com/xmuyulab/htslib).

## 1 Introduction

With the rapid progress of High Throughput Sequencing (HTS) technology, also known as Next Generation Sequencing (NGS) technology [1], the cost of DNA sequencing has been dramatically reduced over the past few years. With reduced costs, NGS technology has found a wider spectrum of applications ranging from basic biomedical research to clinical diagnose, precision medicine, drug discovery, agriculture, forensics, etc. The wide applications of NGS technology has created a huge amount of genomic data, for which efficient and effective representation technology is much needed to handle the corresponding growth in storage costs.

As raw NGS sequencing data are represented as native text files, they can be compressed using general text compression tools, such as gzip (www.gzip.org), bzip2 (www.bzip.org) and 7zip (www.7zip.org). Specific algorithms have also been developed for efficient NGS data compression. Among them, BAM [2] and CRAM [3, 4] are the two most widely adopted algorithms for genome data compression. Other algorithms have also been developed, including QUIP [5], Samcomp [6], DeeZ [7] and LFQC [8]. To leverage the statistical characters of NGS sequencing data for better coding efficiency, most genome data compression algorithms split the sequencing data into separate data streams according to their data types, and compress them individually based on their statistical properties.

Among all data streams from NGS data, per base quality values, which carry information about the likelihood of each base call being in error, is the most challenging component for data compression due to their high entropy. It has been found that quality values can take up to 80% of the lossless compressed file size [9]. On the other hand, when downstream applications are considered, [9] showed that lossy compression of quality values can significantly alleviate the storage while maintaining variant calling performance comparable to that with original data. In some cases, lossy compression can even lead to superior variant calling performance, probably due to the removal of noise in quality values [10].

To improve coding efficiency at lossy rates, many lossy compressors implement highly customized quantization schemes for quality value that either leverage on statistical dependencies among neighbor quality values for rate-distortion optimization encoding [11], or consider downstream analysis to minimize the impact of lossy compression to the analysis results [12, 13, 14]. In QVZ2 [11], quality values are modeled with a Markov chain of order one. Quality values are then quantized based on the Markov model with their positions within the read the previously quantized value as context to maximize the overall rate-distortion performance. NGC [12] pileups the aligned reads with the reference genome, and distinguishes the quality values into four categories depending on the alignment information, e.g., match or mismatch with the reference genome. Different quantization schemas are applied to different categories. CALQ [13] also pileups aligned reads with the reference genome, and further calculates the genotype uncertainty of different genotypes at different loci. Quality values from loci with higher genotype uncertainty are then quantized at smaller quantization step sizes. Crumble [14] used a set of heuristics rules to retain quality values that are essential to downstream analysis, vary by compression level requested. Unnecessary quality values are replaced with constant high values to reduce the entropy for bases that agree with a confident consensus call, or optionally set to a constant low value, heavily quantized, or left intact for bases that disagree with a confident consensus call.

In this paper, we present ScaleQC (Scalable Quality value Compression), a novel quality value compression algorithm that combines lossy and lossless compression in a unified framework. By using a “horizontal” bit-plane scanning and coding approach, ScaleQC generates a compressed quality value bits-stream that can be randomly truncated to any intermediate data rates from lossless to virtually “0” when necessary. The truncated bit-stream can still be decoded and our results show that the quality values decoded from truncated ScaleQC bit-stream can achieve similar or better variant calling accuracy performance when compared to those from other state-of-the-art lossy compressors at the same compressed file sizes. Moreover, the lossless compression performance of ScaleQC is not compromised despite its bit-stream level scalability when compared to other lossless only compressors. To the knowledge of the authors, ScaleQC is the first compressor that provides bit-stream level scalability for lossy to lossless quality value compression. ScaleQC can thus be used as a universal “encode once, decode many times” format of choice for bioinformatic and/or clinic applications where the same set of sequence data are to be used at multiple sites for different applications. ScaleQC has been integrated with SAMtools as an additional quality value coding mode for CRAM. The source code of ScaleQC is available for downloading at (https://github.com/xmuyulab/samtools) and (https://github.com/xmuyulab/htslib).

## 2 Methods

### 2.1 The ScaleQC framework

In this paper, we present ScaleQC for scalable lossy-to-lossless quality value compression for aligned sequencing data. ScaleQC takes SAM/BAM files as inputs and produces scalably compressed quality values that can be further truncated to the desired compression ratio in the future. We integrated ScaleQC with the CRAM tools (https://www.sanger.ac.uk/science/tools/cram) available in SAMtools toolkit, where it can be directly invoked as a special coding mode for quality values. The ScaleQC compressed quality values are stored together with other CRAM data fields in the CRAM file format.

To achieve good data reduction performance, ScaleQC uses adaptive binary arithmetic code [15] that had been successfully used in various video [16] and audio coding algorithms [17], with special context design to capture the statistical properties of quality values. Furthermore, for good variant calling performance at lossy rates, quality values from potential variant sites that are more important for downstream analysis are encoded earlier in the scalable coding process. In such a way, important quality values will be retained when the compressed bit-stream is truncated at higher compression ratios. The framework of ScaleQC is illustrated in Figure 1. The function blocks of ScaleQC are explained in more details in the subsequent sections.

**Figure 1:**
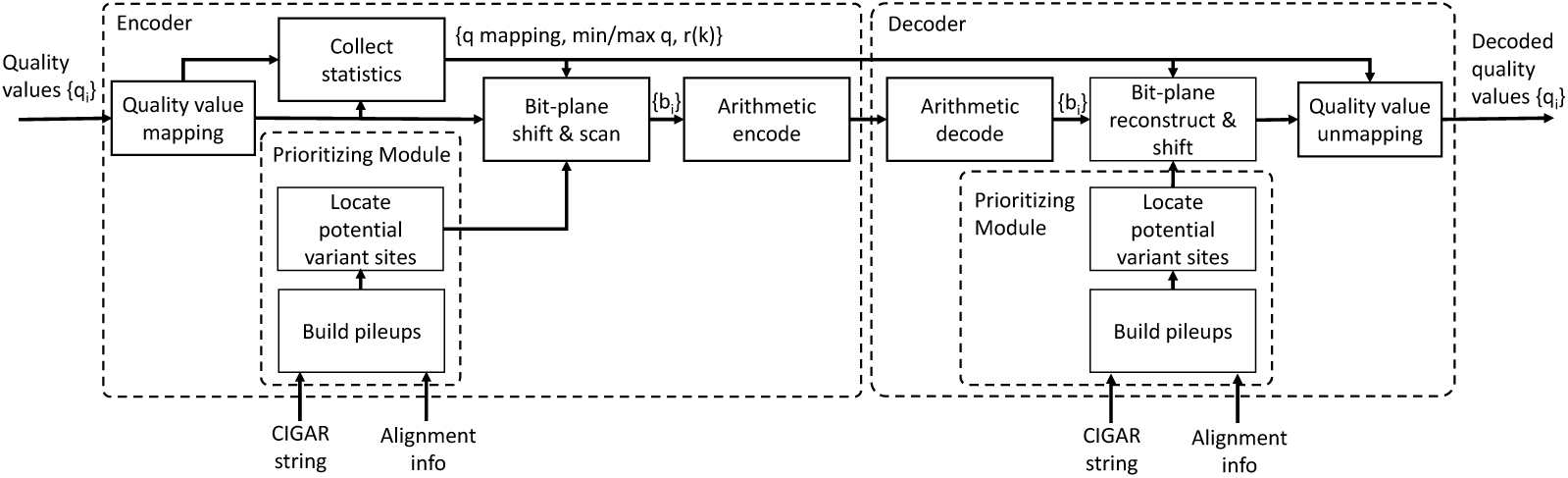
Framework of ScaleQC. Bit-plane encoded quality bits together with side information including quality mapping information, minimum and maximum quality values of different segments, and the context based reconstruction values are transmitted from encoder to decoder to reconstruct the quality values. Partial reconstruction of quality values is possible if only a portion of quality bits are received by the decoder. Potential variant sites are located through pileups of aligned reads that are accessible to both encoder and decoder. Hence, it doesn’t need to be transmitted as side information.

### 2.2 Quality value mapping

Quality values are part of the NGS data stream that indicate the quality of identification of nucleobases generated by DNA sequencing machine. More precisely, quality values are basically integers that indicate the probabilities *p* of the corresponding necleobases in the read sequence being incorrectly identified by the sequencer. They are produced by next generation sequencers during the base call process, and play a significant role in the downstream bioinformatics analysis of NGS data such as variant calling [18].

The most widely used function that converts base call error probability *p* to quality value is the standard Sanger variant for base call reliability assessment calculated as follows:

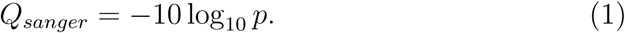

The quality value is further represented as (*Q*_*sanger*_ + 33) using ASCII characters.

Although a wide range of quality values are allowed theoretically (e.g., 33–126), in practice sequencing machines may only use a small subset of them in the output sequencing data. As in bit-plane coding, the number of bits to be processed per encoding symbol is determined by the range of the alphabet set of the symbol (maximum quality value minus minimum quality value), directly performing of bit-plane coding on raw quality values is inefficient. For this reason, we use a lookup table to map the raw quality values to a compact list from 0 to *Q*_*m*_ where *Q*_*m*_ is the total number of valid quality values in the original data. The bit-plane coding is then performed on the quality values after they have been mapped to the compact list. To correctly reconstruct the quality values in the decoder, the lookup table is encoded as a bitmap and sent to the decoder as side information.

### 2.3 Prioritizing of important quality values

The main role of quality values is to help the downstream analysis, in particular, variant caller, to judge whether discrepancies between sequencing data and reference genome are results from true variants or sequencing errors. In this regard, quality values from different loci could carry different levels of importance as those from sites where variant callers may incorrectly give the wrong call are more important compared to other quality values [13, 14].

To maintain variant calling accuracy at lossy rates, ScaleQC builds pileups of aligned reads according to their aligned locations on the reference genome and identifies genome loci with mismatch base calls (including insertion and deletion) as potential variant sites. ScaleQC then amplifies quality values from those potential variant sites by a factor of 2^*s*^, where *s* > 0 is a pre-selected parameter. Equivalently, the amplification lifts the bit-plane of those important quality values by an amount of *s* so that they will be encoded earlier in the bit-plane coding process. In this way, quality values will be decoded to higher fidelity when the scalably compressed bit-stream is truncated to lower data rates. In the decoder, decoded values from those sites are scaled down by the same amount of 2^−*s*^ to restore their original levels. Since the loci of potential variant sites are derived from pileups of aligned reads that are accessible to both encoder and decoder, they don’t need to be transmitted as side information.

### 2.4 Bit-plane coding

To achieve scalable compression, we use bit-plane coding to encode quality values after they have been preprocessed as described earlier. The bit-plane coding process is illustrated in Figure 2, where quality values are grouped and scanned from Most Significant Bit (MSB) to Least Significant Bit (LSB) horizontally to produce the quality bits to be encoded. Without loss of generality, we assume quality value *q* after mapping is bounded by *q*_*min*_ ≤ *q < q*_*max*_, and the binary representation of *q* is then given by

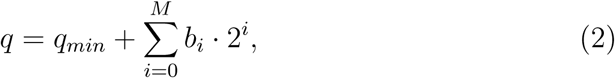

where *M* is the maximum bit-plane, i.e., 2^*M*^ ≥ *q*_*max*_ − *q*_*min*_ > 2^(*M*−1)^, and *b*_*i*_ ∈ {0, 1} denote binary quality bits that represent quality value *q*. The sequential bit-plane coding process is then started from coding of quality bits from highest bit-plane of the whole quality value block after bit-shifting, i.e., from *M*′ = *M* + *s* to LSB 0 as shown in Figure 2. Note that coding of bit-plane at some locations will be skipped if there is no quality bit present after bit-shifting.

**Figure 2:**
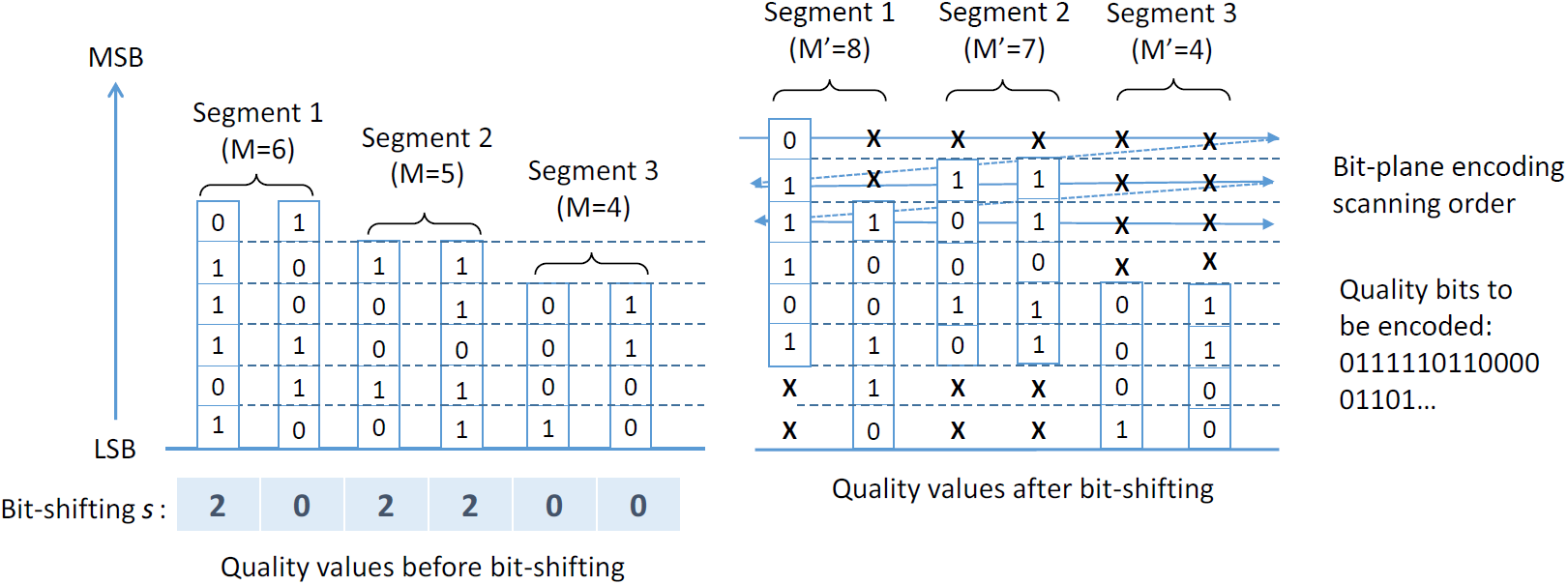
Bit-plane representation and bit-shifting of quality values. MSB for most significant bit, and LSB for least significant bit. During bit-plane encoding, the quality bits will be scanned horizontally from MSB to LSB. *M* indicates original maximum bit-plane for each coding segment and *M*′ = *M* + *s* is the maximum bit-plane after quality values from potential variant sites are shifted by *s* bit-planes. Locations marked with “X” will be skipped during bit-plane scanning as there is no quality bit present after shifting.

In ScaleQC, quality bits generated from the bit-plane scanning process are further encoded using binary arithmetic code. Adaptive modeling is used in the arithmetic coder to learn the empirical distribution of quality bits as they are being compressed so that an increasingly tighter fit to real distribution is achieved on-the-fly. For better coding efficiency, ScaleQC further uses the context coding techniques [19] where a set of contexts are used to deinterleave quality bits into different groups of similar statistical properties, and each group is then encoded with its learned distribution. The details of the context design are given in the next section.

### 2.5 Context design

It is known that the statistical distribution of quality values varies with respect to their positions *t* in the read [20]. For pair-end sequencing, quality values may further depend on whether the read is the first or the second read from the sequencing process. Other than location, the correlation among adjacent quality values can been exploited to improve the coding efficiency of quality values. For bit-plane coding, distributions of quality bits from lower bit-planes could further correlate with those from higher bit-planes when distribution of quality values is not regular [21]. To capture those statistical dependencies for better coding efficiency, we use the following contexts jointly to encode quality bits generated from the bit-plane scanning process.

#### 2.5.1 Coding Segment Context

We split a read into a number of coding segments of fixed length based on location. Assume that segment length is *S*, the coding segment context *C*_*s*_ of a quality value is given by

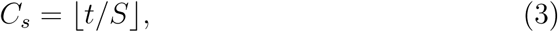

where *t* is its location within the read. For paired-end sequencing reads, we further expand *C*_*s*_ to include whether the quality value is from the first or the second read of a read pair. Since quality values from different coding segments may have very different dynamic ranges, we collect the minimum and maximum quality values for all reads from a coding segment during data preparation. The minimum and maximum quality values of each segment are then used to calculate the starting point of the bit-plane coding process for different coding segments. Note that both values need to be transmitted to the decoder as side information so that the decoder could correctly reconstruct the bit-planes from the decoded quality bits.

#### 2.5.2 Bit-Plane Context

ScaleQC uses bit-plane context defined as *C*_*b*_ = *i* − *M*, where *i* is the bit-plane of the current encoded quality bit, to separate quality bits from different bit-planes into different group as their statistical distribution can be different.

#### 2.5.3 Partial Reconstruction Context

The partial reconstruction context for coding quality value reflects the previously encoded quality bits from higher bit-planes, which is calculated as:

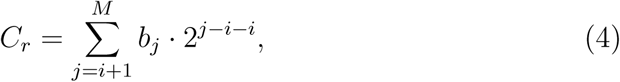

where *i* is the bit-plane of the current encoded quality bit. Note that as *b*_*j*_ (*j* > *i*) will be encoded earlier than *b*_*i*_, the same context can be used for decoding *b*_*i*_ without additional signaling. Partial reconstruction context is particularly useful when the statistical distribution of original quality *q* is rather irregular within its alphabet set.

#### 2.5.4 Adjacent Quality Context

Statistical dependencies among adjacent quality values can be readily exploited by a non-scalable coder such as run-length code [22], entropy coding with a higher-order statistical model [23] or dictionary-based compression [24]. Unfortunately, it is relatively challenging for a scalable coding approach to exploit this dependencies due to limitations from its horizontal bit-plane coding process. In ScaleQC, we only use the quality bit of the same bit-plane from previously encoded quality value as the context for coding the current quality bit. We found that this context provide satisfactory coding efficiency without introducing too much complexity to the bit-plane coding process.

### 2.6 Bitstream structure

The coding and decoding process of ScaleQC is performed on a block by block basis, where each coding block consists of a number of aligned reads, say, 10,000 reads. As in BAM/CRAM design, each coding block of ScaleQC is independent without referring back to previous blocks. Such encoding would produce slighter inferior compression performance due to the overhead in side information signaling and the resetting of the adaptive statistical model in arithmetic codes. However, it allows random access to the records within the compressed files.

The structure of the ScaleQC encoded quality value bit-stream is shown in Figure 3. For each coding block, all the quality values are encoded into a quality data block, which includes a header, a side-information block, followed by the encoded bits for quality values. Note that when necessary, encoded bits portion can be truncated shorter to produce a compressed file of lower data rates without the need to re-encode the whole file.

**Figure 3:**
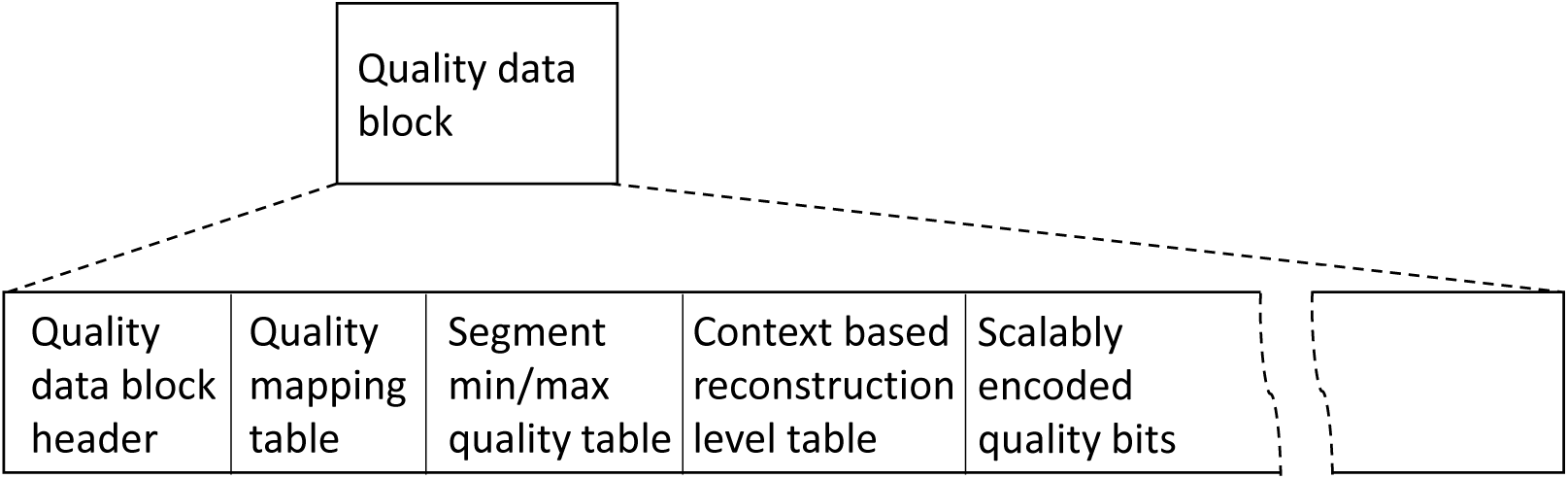
Bit-stream structure for a quality data block in ScaleQC.

### 2.7 Decoding process

Quality value can be losslessly reconstructed at the decoder side by reconstructing the bit-planes as

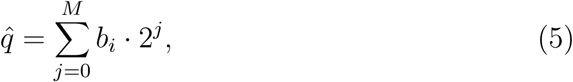

if we have fully received all the quality bits. Moreover, if we only receive partial quality bits from bit-plane *M* to bit-plane *k* (*M* ≥ *k* > 0) at lossy compression rates, it is still possible to recover a partially reconstructed quality value as follows:

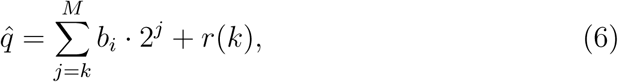

where *r*(*k*) is the reconstruction offset if the bit-plane decoding process is terminated at bit-plane *k. r*(*k*) can be calculated as Minimum Mean Square Error (MMSE) estimation of *q* based on partially reconstructed bit-planes as

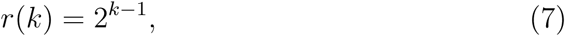

if we assume uniform distribution of *q* within untransmitted bit-planes.

In ScaleQC, to further improve accuracy in quality value reconstruction at lossy compression, we provide an option for the decoder to use context based reconstruction values *r*(*k*), which are calculated at the encoder by averaging the actual quality values for different coding segment and partial reconstructed value contexts and transmitted to the decoder as side information. To reduce the amount of side information, context based *r*(*k*) is used only for larger *k* > *K* where *K* is a user-selectable parameter. For smaller *k*, MMSE reconstruction is still used.

## 3 Results and discussions

To evaluate the performance of ScaleQC, we first compared its lossless compression performance to other lossless quality value compressors. We then studied the impact of lossy compression of quality values on downstream analysis, in particular, variant calling, at different data rates.

### 3.1 Lossless quality value compression of ScaleQC

We tested the compression performance in terms of compression ratio, defined as ratio between the size of compressed quality values and that of original quality values, of ScaleQC on the following four data sets.

- **ERR174324**: Whole Genome Sequencing (WGS) data of NA12878 human sample sequenced by Illumina HiSeq 2000 (http://www.ebi.ac.uk/ena/data/view/ERP001775/). This data was compiled by the Joint Ad-hoc Group on Genomic Information Compression and Storage (JAhG) between ISO/IEC JTC 1/SC 29/WG 11, also known as Moving Picture Experts Group (MPEG), and ISO/TC 276/WG 5 for the development of MPEG standardization of genomic information representation. We selected the first pair of fastq files from dataset H01, namely ERR174324_1.fastq and ERR174324_2.fastq, which has a coverage of approximate 14x.
- **Readshift**: WGS data of of NA12878 human sample sequenced using Illumina Novaseq from Readshift project (https://github.com/dnanexus/Readshift). We selected one pair of fastq files from the dataset, NA12878_I30_S4_L001, which can be downloaded from https://platform.dnanexus.com/projects/F9K5zYQ0ZQBb5jJf6X2zZPv1/data/. This data has an approximate coverage of 30x.
- **Garvan**: WGS data of NA12878 human sample sequenced using Illumina HiSeq X by Garvan Institute. Available from PrecisionFDA Consistency Challenge (https://precision.fda.gov/challenges/consistency). This data has an approximate coverage of 50x.
- **HG002**: WGS data of HG002 (NA24385) human sample sequenced using Illumina HiSeq 2500 by NIST, available from PrecisionFDA Truth Challenge (https://precision.fda.gov/challenges/truth). This data has an approximate coverage of 50x.

In order to have a fair comparison, we used only lossless compression tools that provide random access in our evaluation, which include CRAM v3 [4], DeeZ [7] and TSC [25]. The comparison results are summarized in Table 1. It can be seen that lossy to lossless scalability is achieved by ScaleQC with very marginal overheads in lossless compression. The highest overhead is at around 10% for low coverage data (ERR174324, 14x) when compared to the best performer (DeeZ). In some cases, ScaleQC can even outperform lossless-only compressors. For example, for Novaseq data ScaleQC achieves better compression performance compared to other lossless compressors.

**Table 1:** Comparison of compressed file sizes (MB), compressed quality value sizes (MB) and compression ratio (CR) for different lossless compressors.

| Data | Method | FileSize(MB) | QualSize(MB) | CR |
| --- | --- | --- | --- | --- |
| ERR174324<br>(HiSeq 2000, 14x) | CRAM | 336.4263 | 269.1801 | 0.3164 |
|  | DeeZ | 324.7912 | 267.0748 | 0.3139 |
|  | ScaleQC | 366.9106 | 299.6628 | 0.3522 |
|  | TSC | 479.0398 | 333.4339 | 0.3919 |
| Readshift<br>(Novaseq, 30x) | CRAM | 214.8812 | 95.7802 | 0.0579 |
|  | DeeZ | 212.8942 | 103.2330 | 0.0624 |
|  | ScaleQC | 205.2750 | 86.1840 | 0.0521 |
|  | TSC | 389.9121 | 155.4523 | 0.0940 |
| Garvan<br>(HiSeq X, 50x) | CRAM | 585.4211 | 387.4136 | 0.1475 |
|  | DeeZ | 578.6992 | 392.2624 | 0.1498 |
|  | ScaleQC | 609.0370 | 410.7252 | 0.1564 |
|  | TSC | 896.6827 | 530.5268 | 0.2020 |
| HG002<br>(HiSeq 2500, 50x) | CRAM | 1088.6990 | 881.2713 | 0.3128 |
|  | DeeZ | 1052.7469 | 880.5957 | 0.3125 |
|  | ScaleQC | 1183.3113 | 975.8583 | 0.3463 |
|  | TSC | 1461.4529 | 1097.5805 | 0.3895 |

### 3.2 Impact of lossy quality value compression on variants calling

We used the widely adopted GATK pipeline as our variant calling pipeline to study the impact of lossy quality value compression. Briefly, raw sequencing reads were mapped to the UCSC hg19 reference sequence with BWA [2]. PCR duplicates were removed by Picard. Variants were detected using GATK HaplotypeCaller (HC) [26]. Indel realignment was not performed as it is not recommended for GATK after version 3.6. In addition, we omitted Base Quality Score Recalibration (BQSR) in our evaluation pipeline since it may obscure the impact of lossy compression on final variant discovery results. Raw VCF output produced from GATK HC were filtered using hard filtration using parameters recommended on GATK website. The final variant calling results were analyzed using the hap.py benchmarking tools as proposed by Illumina and adapted by the Global Alliance for Genomics and health (GA4GH, https://github.com/ga4gh/benchmarking-tools). We refer to the Supplementary Material for specific command lines and options of all tools used in our evaluation.

We used Recall (*R*), Precision (*P*), and F1 (*F*_1_) scores defined below as our comparison metrics:

- Recall: Proportion of called variants that are included in the ground-truth set, i.e., *R* = *TP/*(*TP* + *FN*),
- Precision: Proportion of ground-truth variants within all the variants called by the variant calling pipeline, i.e., *P* = *TP/*(*TP* + *FP*),
- F1 score: Harmonic mean of precision and recall defined as *F*_1_ = 2 · *P · R/*(*P* · + *R*)

Here, *TP* (True Positive) is defined as the number of variants that are in both the ground-truth set and set of called variants; *FP* (False Positives) is defined as the number of variants that are in the called set but not in the ground-truth set; *FN* (False Negative) is defined as the number of variants that are in the ground-truth set, but not in the set of called variants.

We used the same data sets as in the previous section for our study on lossy quality compression. As the primary goal of our study is to identify the impact of lossy compression on the downstream analysis, we didn’t use the consensus sets of variants released together with these datasets as our ground-truth sets for comparison. Instead, we use the variants called from original sequencing data using our variant calling pipeline as our ground-truth sets for comparison. Since those consensus sets are typically collections of high confident variants discovered from sources other than the provided data [27], directly comparing called variants from lossy compressed data to those from original data could better reflect the effects of lossy quality value compression on the downstream analysis pipelines.

We compared ScaleQC to eight lossy compressors, including Illumina binning (performed with DSRC) [28], P/R-Block [29], Quartz [10], LEON [30], QVZ2 [11], CALQ [13] and Crumble [14]. Four of the eight tools (P-block, R-block, QVZ2, Crumble) provide variable compression rates by adjusting parameters, while the other four (CALQ, LEON, Quartz, DSRC) have only one fixed rate-distortion point without tunable parameters. ScaleQC provides bit-stream level scalability from nearly 0 (when all quality values are discarded and only the header information is retained) to lossless. Fig. 4 compares the F1 score with respect to SNP (Single Nucleotide Polymorphism) and INDEL (Insertion and Deletion) of different lossy compressors on the four sequencing data under evaluation. Detailed results in terms of Recall and Precision are provided in the Supplementary Material.

**Figure 4:**
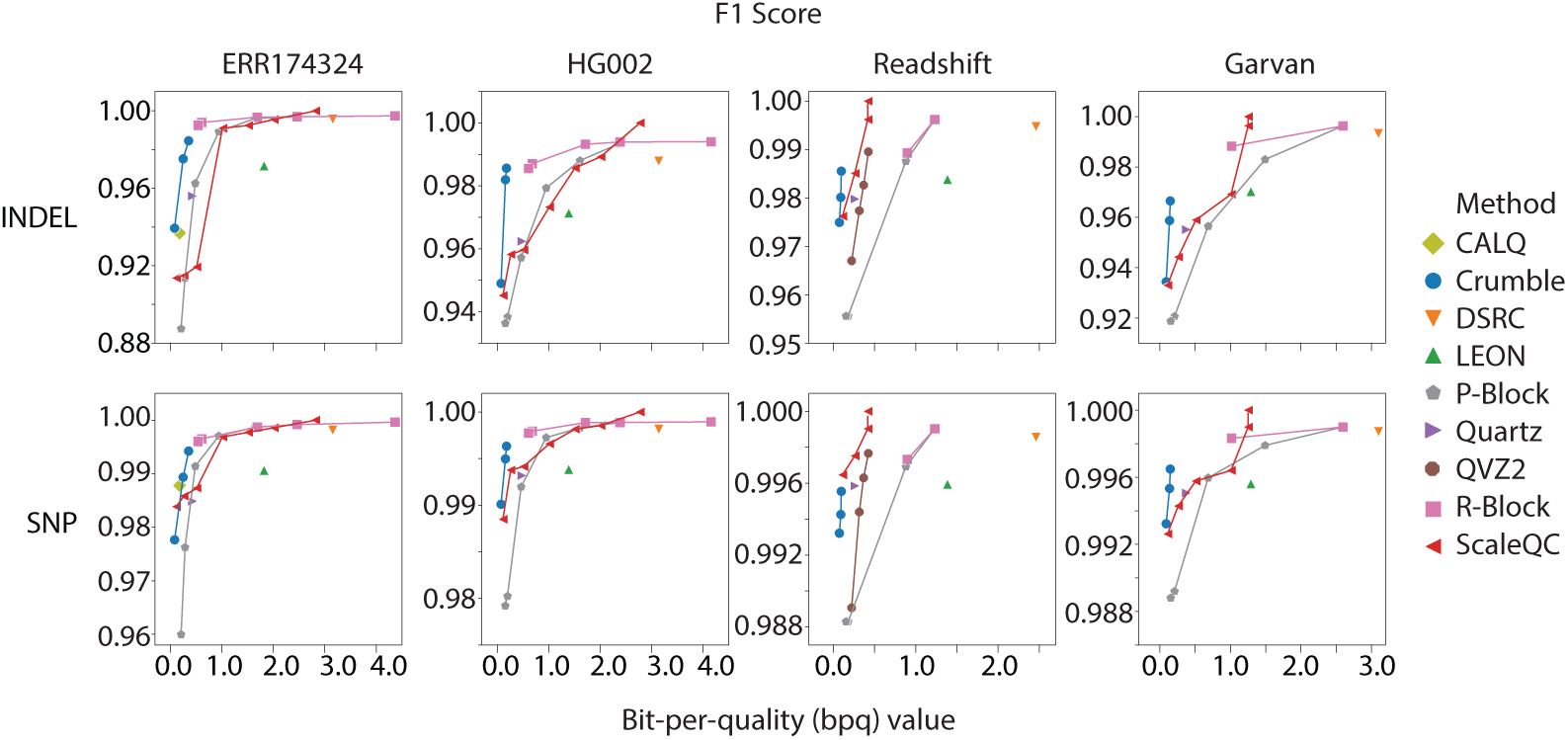
F1 score of variant calling results when quality values are compressed using different lossy compressors. For ScaleQC, the highest data points are corresponding to lossless coding (F1=1). Missing data points of some compressors are caused by either crashes or compatibility issues.

As expected, higher coverage data (Garvan and HG002) shows less sensitivity to changes from lossy compression of quality values while performance on lower coverage data (ERR174324) degrades most when quality values are heavily compressed. Lossy compressors that simply use quality value binning without considering downstream analysis, such as Illumina binning, provide only mediocre compression performance. On the other hand, algorithms that prioritize quality values based on downstream analysis, such as Crumble and CALQ, among others, perform very well at low data rates. Regarding the proposed algorithm, ScaleQC achieves similar accuracy performance compared to other lossy compressors although its bit-stream is obtained through truncation from lossless compressed bit-stream without re-encoding. In fact, for Novaseq data, it outperforms other lossy compressors at all data rates under evaluation.

### 3.3 Computational performance

The run-time of all tests are recorded as in Table 2 for all lossy and lossless compression tools under evaluation. For tools supporting multi-threading, we made all the 48 threads available on the testing platform to the software under evaluation and set the number of threads to 48 through command line option if available. Note that some tools (CALQ, QVZ2, TSC, P-block, R-block) only support single-thread mode, while others (LEON, DSRC, Quartz, DeeZ, CRAM, ScaleQC) support multi-threading. Crumble supports “partial” multi-threading for the BGZF compression component when writing the BAM file. Even for tools supporting multi-threading, the performance on 48 threads shall not be regarded as a linear speedup of the performance on a single thread due to the small file size and the overheads and programming constraints in multi-threading implementations. For example, we noticed that with DeeZ, the runtime for compression and decompression using 48 threads shows only marginal improvements over using 4 threads. Hence, the results presenting here only reflect the maximum possible speed of the software on the testing platform, and cannot be regarded as a benchmark for the numeric complexity of the underlying algorithms.

**Table 2:** Computing time for quality value compression and decompression for lossless and lossy compressors under evaluation. For lossy compressor, computing time at different command line options are given. Since ScaleQC provides bit-stream scalability, compression time is given only at lossless rate. NA indicates missing data point caused by either crashes or compatibility issues.

| Method | Param | Garvan (HiSeq X, 50x) |  | ERR174324 (HiSeq 2000, 14x) |  | Readshift (Novaseq, 30x) |  | HG002 (HiSeq 2500, 50x) |  |
| --- | --- | --- | --- | --- | --- | --- | --- | --- | --- |
|  |  | Comp | Decomp | Comp | Decomp | Comp | Decomp | Comp | Decomp |
| CRAM | lossless | 0m25.717s | 0m12.447s | 0m32.135s | 0m29.919s | 0m28.040s | 0m16.494s | 0m34.020s | 0m33.989s |
| DeeZ | lossless | 0m53.193s | 0m35.319s | 2m6.494s | 1m32.657s | 1m10.448s | 0m45.386s | 2m15.432s | 1m29.791s |
| TSC | lossless | 3m46.062s | 2m24.682s | 9m48.937s | 6m28.492s | 5m31.204s | 4m10.295s | 10m38.726s | 6m57.814s |
| CALQ |  | NA | NA | 26m35.129s | 5m6.511s | 45m34.182s | 9m48.247s | 78m19.648s | 16m38.096s |
| DSRC |  | 0m28.926s | 0m14.817s | 0m14.932s | 0m19.330s | 0m18.192s | 0m12.915s | 0m10.971s | 0m14.548s |
| LEON |  | 1m44.505s | 0m50.261s | 0m46.103s | 0m21.076s | 1m11.804s | 0m42.472s | 1m50.018s | 1m1.985s |
| Quartz |  | 13m58.658s | NA | 4m58.977s | NA | 11m20.246s | NA | 14m21.593s | NA |
| Crumble | 1 | 4m42.552s | NA | 2m4.256s | NA | 3m8.154s | NA | 4m52.245s | NA |
|  | 5 | 3m58.909s | NA | 1m17.422s | NA | 2m19.337s | NA | 4m5.815s | NA |
|  | 9 | 4m2.226s | NA | 1m13.015s | NA | 2m20.291s | NA | 4m8.483s | NA |
| QVZ2 | 1 | NA | NA | NA | NA | 4m34.003s | 1m17.266s | NA | NA |
|  | 4 | NA | NA | NA | NA | 4m32.385s | 1m17.046s | NA | NA |
|  | 8 | NA | NA | NA | NA | 4m30.803s | 1m13.354s | NA | NA |
|  | 16 | NA | NA | NA | NA | 4m32.396s | 1m13.120s | NA | NA |
| P-block | 1 | 1m14.541s | 0m25.711s | 0m28.434s | 0m9.188s | 0m38.524s | 0m12.504s | 1m23.735s | 0m27.236s |
|  | 2 | 1m11.599s | 0m36.612s | 0m24.032s | 0m11.022s | 0m36.733s | 0m12.378s | 1m12.122s | 0m22.284s |
|  | 4 | 1m2.949s | 0m31.042s | 0m23.387s | 0m8.931s | 0m37.366s | 0m12.230s | 1m0.661s | 0m20.158s |
|  | 8 | 0m50.651s | 0m24.001s | 0m19.416s | 0m7.331s | 0m33.782s | 0m10.705s | 0m51.037s | 0m11.849s |
|  | 16 | 0m41.430s | 0m17.954s | 0m17.615s | 0m6.608s | 0m28.797s | 0m5.757s | 0m45.888s | 0m7.901s |
|  | 32 | 0m41.658s | 0m16.028s | 0m16.937s | 0m5.339s | 0m27.929s | 0m4.514s | 0m45.047s | 0m5.562s |
| R-block | 1.05 | 1m18.986s | 0m26.303s | 0m42.518s | 0m11.881s | 0m40.214s | 0m12.300s | 1m53.970s | 0m39.222s |
|  | 2.6 | 1m19.666s | 0m38.062s | 0m36.174s | 0m11.885s | 0m40.320s | 0m12.314s | 1m50.699s | 0m35.193s |
|  | 4.2 | 1m20.502s | 0m37.219s | 0m31.441s | 0m9.370s | 0m38.088s | 0m12.412s | 1m30.760s | 0m27.146s |
|  | 7.4 | 1m20.152s | 0m28.107s | 0m25.792s | 0m7.788s | 0m39.697s | 0m12.244s | 1m20.002s | 0m22.822s |
|  | 25 | 0m58.405s | 0m26.959s | 0m18.631s | 0m4.833s | 0m38.401s | 0m12.494s | 1m0.656s | 0m20.191s |
|  | 30 | 0m55.452s | 0m21.411s | 0m18.048s | 0m4.524s | 0m35.594s | 0m10.814s | 0m56.426s | 0m12.655s |
| ScaleQC | 0.1 |  | 1m24.716s |  | 0m34.919s |  | 1m5.445s |  | 1m32.228s |
|  | 0.25 |  | 1m25.665s |  | 0m33.769s |  | 1m7.446s |  | 1m32.195s |
|  | 0.5 |  | 1m22.499s |  | 0m33.205s |  | 1m9.820s |  | 1m31.037s |
|  | 1 |  | 1m20.068s |  | 0m33.070s |  | 1m5.682s |  | 1m31.362s |
|  | 1.5 |  | 1m24.123s |  | 0m31.877s |  | 1m3.457s |  | 1m31.278s |
|  | 2 |  | 1m20.189s |  | 0m32.686s |  | 1m1.442s |  | 1m33.552s |
|  | lossless | 1m24.984s | 1m23.761s | 0m48.730s | 0m33.024s | 1m3.796s | 1m4.040s | 1m40.130s | 1m33.848s |

Secondly, some tools only parse and compress/decompress the quality strings in pre-extracted quality files (QVZ2) or in FASTQ files (LEON, DSRC, Quartz), some others parse SAM/BAM files and compress/decompress only the quality strings (CALQ, P-block, R-block), while others (CRAM, ScaleQC) provide the full functionality to parse the entire SAM/BAM file and compress/decompress all the fields in the SAM file. Crumble parse SAM/BAM files and also output in BAM format. Although compression and decompression of quality values could be the most time-consuming components of the entire process, tools that provide full functionality could still be slowed down by the overheads in parsing and compression of read names, nucleotide bases, etc.

Despite the above limitations in computational performance comparison, we still can see that ScaleQC provides a complexity performance that is on par with other compressors. Overall, the computing time of ScaleQC, in particular, the decoding time is still higher than that fastest lossy or lossless compressors under evaluation (DSRC, CRAM), but difference is not outrageous. The main computational overhead of ScaleQC is due to its bit-plane coding process to provide the desired scalability. As ScaleQC is a new tool, we envision that its complexity could be further reduced through implementation and/or algorithm optimization in the future. For example, it is possible to integrate other lossy compressors, such as Crumble, as a core coder for ScaleQC such that ScaleQC only encodes the difference between the original and the decoded quality value from the core compressor. In this way, not only the computational complexity can be reduced due to the reduced range of scalable coding, but also the performance at extremely low data rates can be improved.

### 3.4 Impacts of lossy quality values compression on overall file sizes

As quality values occupy a significant portion of the total sequencing data file, the overall compressed file sizes highly related to the sizes of quality values after compression (Fig. 5). At the highest bit-per-quality (bpq) values, ScaleQC achieves lossless compression of quality values and helps to reduce the overall compressed file size to 0.35-0.6 of its original BAM file sizes. The highest lossless compression ratio is around 0.35 for Novaseq data. Interestingly, despite their poor compressibility at lossless rates, two HiSeq 2000 and HiSeq 2500 data (HG002, ERR174324) achieve a greater amount of file size reduction when the quality values are lossy compressed. At bpq=0.2, ScaleQC can reduce the size of sequence files to 0.1-0.25 of its original sizes in BAM format, which is significant. It is possible to further reduce the overall file size by further truncating the ScaleQC quality bit-stream. However, the impact on the accuracy of downstream analysis has to be considered to determine the best trade-off between storage space saving and analysis quality.

**Figure 5:**
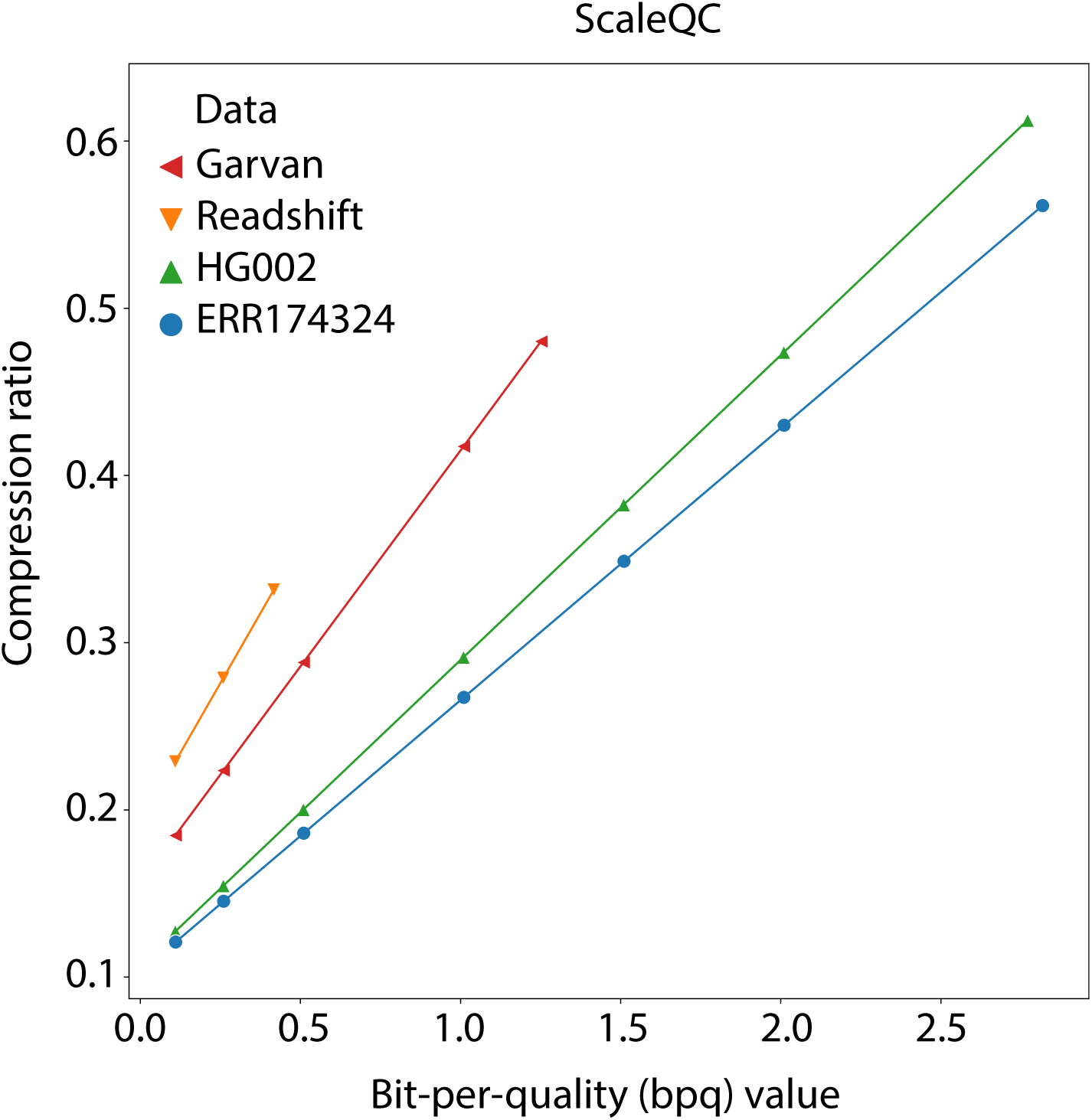
Overall compression ratio of ScaleQC with respect to file sizes of original BAM files at different bit-per-quality values for different sequencing files. As quality values occupy a significant portion of total sequencing data file, the overall compression ratio is highly related to the bpq. Lossless compression is achieved at highest bpq.

## 4 Conclusion

We present ScaleQC, the first quality value compressor that provides bit-stream level lossy to lossless scalability. ScaleQC bridges the gap between lossy and lossless compression of NGS sequencing data, and provides an “encoding once, decoding many time” solution to cater for the needs from applications with different quality fidelity requirements.

Besides scalability, ScaleQC still achieves a good balance among functionality, complexity, and coding efficiency performance. For lossy compression, ScaleQC performs as well as or even better than many state-of-the-art lossy compressors in terms of recall and precision on datasets under test. At the same time, for lossless compression, ScaleQC achieves a size reduction on par with the best lossless quality value compressor available nowadays.

## Acknowledgements

We would like to thank Shun Wang for help in testing and evaluating the program, and Canqiang Xu for visualization of the results.

