## Supplementary for "ScaleQC: A Scalable Lossy to Lossless Solution for NGS Sequencing Data Compression"

Rongshan Yu and Wenxian Yang

### 1 Tools for benchmarking

#### 1.1 Lossy compression

We compared ScaleQC with eight compression tools that aim at lossy compression of base quality values. The tools for benchmarking with specific version numbers used in our experiments are listed below:

- CALQ [1], version 1.0.0, 194fdea, built from revised version release for bioinformatics, (`https://github.com/voges/calq`)
- P-block, R-block [2], da36a12, built from source, (`https://github.com/rcanovas/libCSAM.git`)
- QVZ2 [3], 70e5926, built from source, (`https://github.com/mikelhernaez/qvz2.git`)
- LEON [4], version 1.0.0, download built executables, (`http://gatb.inria.fr/software/leon/`)
- Crumble [5], version 0.8.3, 3fbaaaa, built from source, (`https://github.com/jkbonfield/crumble.git`)
- Quartz [6], version 0.2.2, e843195, built from source, (`https://github.com/yunwilliamyu/quartz.git`)
- DSRC [7] (implements IlluminaBinning (`http://res.illumina.com/documents/products/whitepapers/whitepaper_datacompression.pdf`), version 2.00 RC @ 28.03.2014, executables downloaded from (`http://sun.aei.polsl.pl/REFRESH/index.php?page=projects&project=dsrc&subpage=download`)
- ScaleQC, our method, version 1.0, available from (`https://github.com/xmuyulab/samtools`) and (`https://github.com/xmuyulab/htslib`)

The inputs of these tools include various file types such as .bam (Crumble, ScaleQC) or .sam (CALQ, P-block, R-block) file, .fastq file (LEON, QUARTZ, DSRC), and .qual file (QVZ2). Here, .qual files are text files containing only the base quality strings, one line per base quality string. Therefore, the content of a .qual file is identical as every fourth line of its corresponding .fastq file.

Correspondingly, the outputs of these tools include different file types. Crumble outputs BAM files and we further compress these BAM files into .cram files using CRAM (by `samtools view`). ScaleQC outputs .sqc file that is compatible with CRAM. The sizes of compressed quality values in .cram and .sqc files can be then obtained by `cram_size`. Quartz outputs .fastq files with replaced base quality values. We extracted the decoded quality values into .qual file. As recommended by the author ( [6]), we then used `bzip2` with default parameters to further compress the .qual files into .qual.bz2 files and the bitstream size is evaluated by `wc -c file.qual.bz2`. Other tools output a bitstream file containing only the compressed quality values in their own format. CALQ outputs .cq files, DSRC outputs .dsrc files, LEON outputs .leon files, P-block and R-block output .cqual files, and QVZ2 outputs .qvz2 files. The size for these files can be obtained by `wc -c file`. These bitstream files are then decompressed by each individual tool to restore the quality values either in .qual or in .fastq files, and then fed back to .sam files by `replace_qual_sam.py` for further variant calling and evaluation.

#### 1.2 Lossless compression

Among the available tools, only CRAM, DeeZ and TSC provide random-access. Therefore we compared ScaleQC with these three tools for lossless compression.

- CRAM v3 [8], as implemented in SAMtools package version 1.9-dirty, (`https://github.com/samtools/samtools.git`)
- DeeZ [9], git commit id 92cd56b, built from source (`https://github.com/sfu-compbio/deez.git`)
- TSC [10], git commit id ebbe931, built from source (`https://github.com/voges/tsc.git`)

The sizes of compressed quality values by DeeZ and TSC can be retrieved from the output log when verbose mode is enabled. For CRAM and ScaleQC, the sizes of compressed quality values can be retrieved from the compressed files using `cram_size`.

#### 1.3 Auxiliary tools

Besides, a number of auxiliary software tools or libraries were installed as dependency of the compression tools, or for evaluation purpose. The source and version of there tools are described as follows.

- BWA, version: 0.7.17-r1198-dirty, (`https://github.com/lh3/bwa.git`)
- GATK, version: 4.1.3 (with HTSJDK version: 2.20.1 and Picard version: 2.20.5), (`https://software.broadinstitute.org/gatk/download/`)
- HTSlib, version: 1.9-dirty, (`https://github.com/samtools/htslib.git`)
- SAMtools, version: 1.9-dirty, (`https://github.com/samtools/samtools.git`)
- replace_qual_sam.py, as provided by CALQ package
- cram_size, as provided by Staden package which is directly installed from Ubuntu 16.04 repository
- hap.py and som.py, built from (`https://github.com/Illumina/hap.py`)

#### 1.4 Detailed description of tools usage

Command lines for compression/decompression of quality values and measuring the size of compressed quality values for each tool used in our benchmarking experiments are listed as follows.

##### 1.4.1 CALQ

```
time calq -r ${ref} ${in}.sam -o ${in}.cq
time calq -d -s ${in}.sam in.cq -o ${in}.cq.sam
replace_qual_sam.py ${in}.sam in.${in}.sam 1> calq.sam
wc -c ${in}.cq
```

##### 1.4.2 P-block and R-block

**P-block**

```
# we tested with pvalues of {1,2,4,8,16,32}
time CompressQual ${in}.sam -q 1 -l <pvalue>
time DecompressQual ${in}.sam.cqual
replace_qual_sam.py ${in}.sam ${in}.sam.cqual.qual 1> pblock.sam
wc -c ${in}.sam.cqual
```

**R-block**

```
# we tested with rvalues of {1.05, 2.6, 4.2, 7.4, 25, 33}
time CompressQual ${in}.sam -q 2 -l <rvalue>
time DecompressQual ${in}.sam.cqual
replace_qual_sam.py ${in}.sam ${in}.sam.cqual.qual 1> rblock.sam
wc -c ${in}.sam.cqual
```

##### 1.4.3 QVZ2

```
# we tested with tvalues of {1, 4, 8, 16}
time qvz2 -t <tvalue> -v -u ${in}.qvz2.qual ${in}.qual ${in}.qvz2
replace_qual_sam.py ${in}.sam ${in}.qvz2.qual 1> qvz2.sam
wc -c ${in}.qvz2
```

##### 1.4.4 LEON

```
time leon -file ${in}.fastq -c
time leon -file ${in}.fastq.leon -d
awk 'NR%4==0 {print}' ${in}.fastq.d > ${in}.fastq.d.qual
replace_qual_sam.py ${in}.sam ${in}.fastq.d.qual 1> leon.sam
wc -c ${in}.fastq.leon
```

##### 1.4.5 Crumble

```
# we tested with vvalues of {1,5,9}
time crumble -v -<vvalue> ${in}.bam ${in}.crumble.bam -O bam,nthreads=48
samtools view -C ${in}.crumble.bam -o ${in}.crumble.cram -T ${ref}
cram_size ${in}.crumble.cram | grep QS
```

##### 1.4.6 Quartz

```
time quartz dec200.bin.sorted 50 48 1 ${in}.fastq
awk 'NR%4==0' ${in}.fastq.filtered_50 > ${in}.quartz.qual
replace_qual_sam.py ${in}.sam ${in}.quartz.qual > ${in}.quartz.sam
bzip2 ${in}.quartz.qual
wc -c ${in}.quartz.qual.bz2
```

##### 1.4.7 DSRC

```
time dsrc c -t48 -l -m2 ${in}.fastq ${in}.dsrc
time dsrc d -t48 ${in}.dsrc ${in}.dsrc.fastq
awk 'NR%4==0 {print}' ${in}.dsrc.fastq > ${in}.dsrc.qual
replace_qual_sam.py ${in}.sam ${in}.dsrc.qual 1> dsrc.sam
wc -c ${in}.dsrc
```

##### 1.4.8 DeeZ

```
time deez -v 2 -t 48 -r ${ref} ${in}.sam -o {in}.dz
time deez -t 48 -r ${ref} ${in}.dz -o ${in}.dz.sam
```

##### 1.4.9 TSC

```
time /mnt/md0/tools/tsc/build/bin/tsc -s ../data/chr.sam -o chr.tsc
time /mnt/md0/tools/tsc/build/bin/tsc -d -s chr.tsc -o chr.tsc.sam
```

##### 1.4.10 ScaleQC

Note that ScaleQC approach has been integrated into SAMtools and HTSlib, therefore the executable for ScaleQC is `samtools` which is compiled from the integrated code.

```
# we tested with bpq values of {0.1, 0.25, 0.5, 1, 1.5, 2, 8}
# encode with lossless mode of bpq 8, and decode with different bpq values
time samtools view -C ${in}.bam -o ${in}.sqc -T ${ref} --sqc-enc -p 8 -@48
time samtools view ${in}.sqc -h -T ${ref} -o ${in}.sqc.bam -p ${bpq} --sqc-dec -@48
cram_size ${in}.sqc | grep QS
```

#### 2 Alignment, variant calling and evaluation

In our experiments, the raw .sam file output from BWA MEM aligner was sorted and indexed by SAMtools, and aligned reads for chromosome 20 were extracted by SAMtools for benchmarking. Reads (.fastq) and quality files (.qual) were converted from .bam file. Note that CALQ and ScaleQC require the input SAM/BAM to be coordinate-sorted, as to build the pile up of SAM records.

For all experiments, we used human reference genome `hs37d5.fa` for alignment. It can be downloaded from (`ftp://ftp-trace.ncbi.nih.gov/1000genomes/ftp/technical/reference/phase2_reference_assembly_sequence/hs37d5.fa.gz`). The command lines to prepare the input files are as follows:

```
bwa mem -t 48 -M $ref ${in}_r1.fastq.gz ${in}_r2.fastq.gz > ${in}.sam
# sam to sorted bam
samtools sort -@48 -h -b ${in}.sam -o ${in}.bam
samtools index ${in}.bam
# extract chromosome 20
samtools view -h -b ${in}.bam chr20 > ${in}_chr20.bam
# extract fastq from sorted bam
samtools view ${in}_chr20.bam | awk '{print "@"$1"\n"$10"\n+\n"$11}' > ${in}_chr20.fastq
# extract quality values from fastq
awk 'NR%4==0 {print}' ${in}_chr20.fastq > ${in}_chr20.qual
```

In order to assess the effect on lossy compression of quality value on the downstream variant calling process, we followed the GATK best practices for germline variant calling using HaplotypeCaller with hard filtering. The evaluation was done by comparing the called variants on decoded sequencing data with those on original BAM file using *hap.py*. The detailed command lines are listed as follows.

```
java -jar $PICARD MarkDuplicates I=${in}.bam O=${in}_mdup.bam M=metrics.txt ASSUME_SORTED=true

$GATK --java-options \"-Xmx20G -Djava.io.tmpdir=./\" HaplotypeCaller -R $ref \
-I ${in}_mdup.bam -L $interval -O ${in}.vcf.gz"

# subset to SNPs-only and indels-only callsets
$GATK SelectVariants -V ${in}.vcf.gz -select-type SNP -O ${in}_snps.vcf.gz
$GATK SelectVariants -V ${in}.vcf.gz -select-type INDEL -O ${in}_indels.vcf.gz

# hard filtering
$GATK VariantFiltration -V ${in}_snps.vcf.gz \
-filter "QUAL < 30.0" --filter-name "QUAL30" \
-filter "SOR > 3.0" --filter-name "SOR3" \
-filter "FS > 60.0" --filter-name "FS60" \
-O ${in}_snps_filtered.vcf.gz

$GATK VariantFiltration -V ${in}_indels.vcf.gz \
-filter "QUAL < 30.0" --filter-name "QUAL30" \
-filter "FS > 200.0" --filter-name "FS200" \
-O ${in}_indels_filtered.vcf.gz

# merge filtered snp and indels into final vcf
$GATK SortVcf -I ${in}_snps_filtered.vcf.gz -O ${in}_snps_sort.vcf.gz
$GATK SortVcf -I ${in}_indels_filtered.vcf.gz -O ${in}_indels_sort.vcf.gz
$GATK MergeVcfs -I ${in}_snps_sort.vcf.gz -I ${in}_indels_sort.vcf.gz -O ${in}_final.vcf.gz

python hap.py ${TRUEVCF} ${in}_final.vcf.gz -o eval_${in} -r ${ref} -V
```

We follow the recommendations from (`https://software.broadinstitute.org/gatk/documentation/article?id=23216`) for hard filtering of the raw variants.

### 3 Testing platform

All experiments are carried out on Ubuntu 16.04, Dual Intel(R) Xeon(R) CPU E5-2678 v3 @ 2.50GHz, 48 threads, 256GB memory.

### 4 Additional results on impacts of lossy quality compression on variant calling

Figure 1 shows the comparison of performance of different lossy compression tools, with regards to the precision, recall and F1-score for variants called on decompressed sequencing data at different compression ratios.

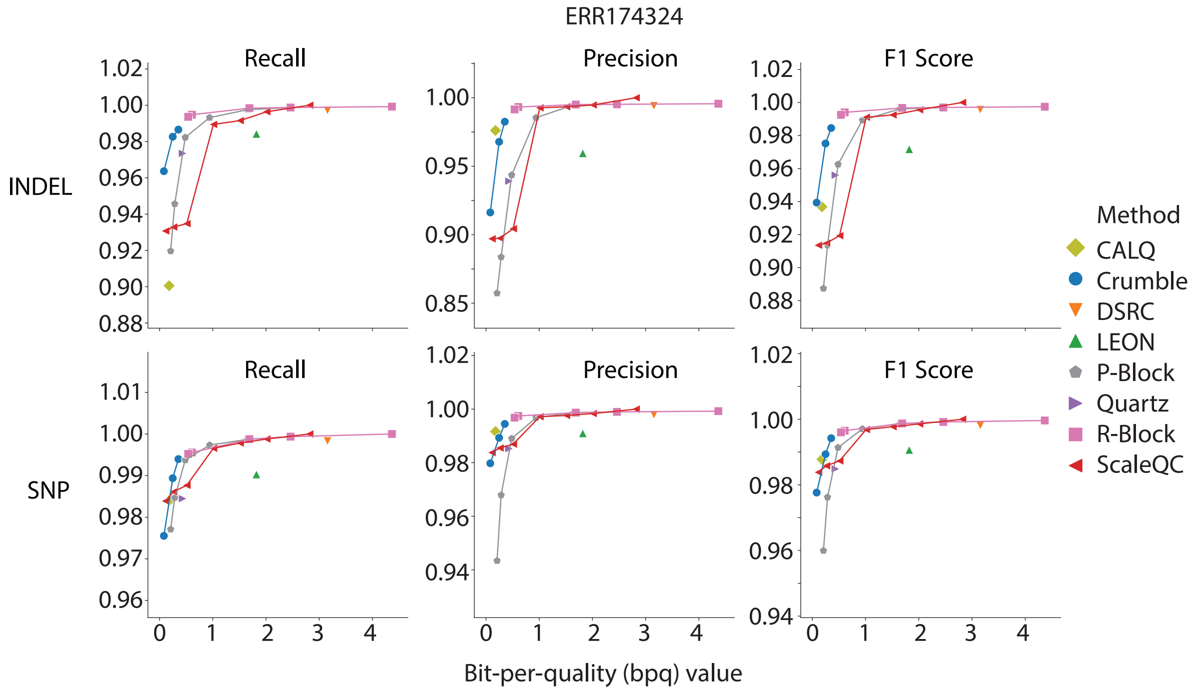


(a) ERR174324

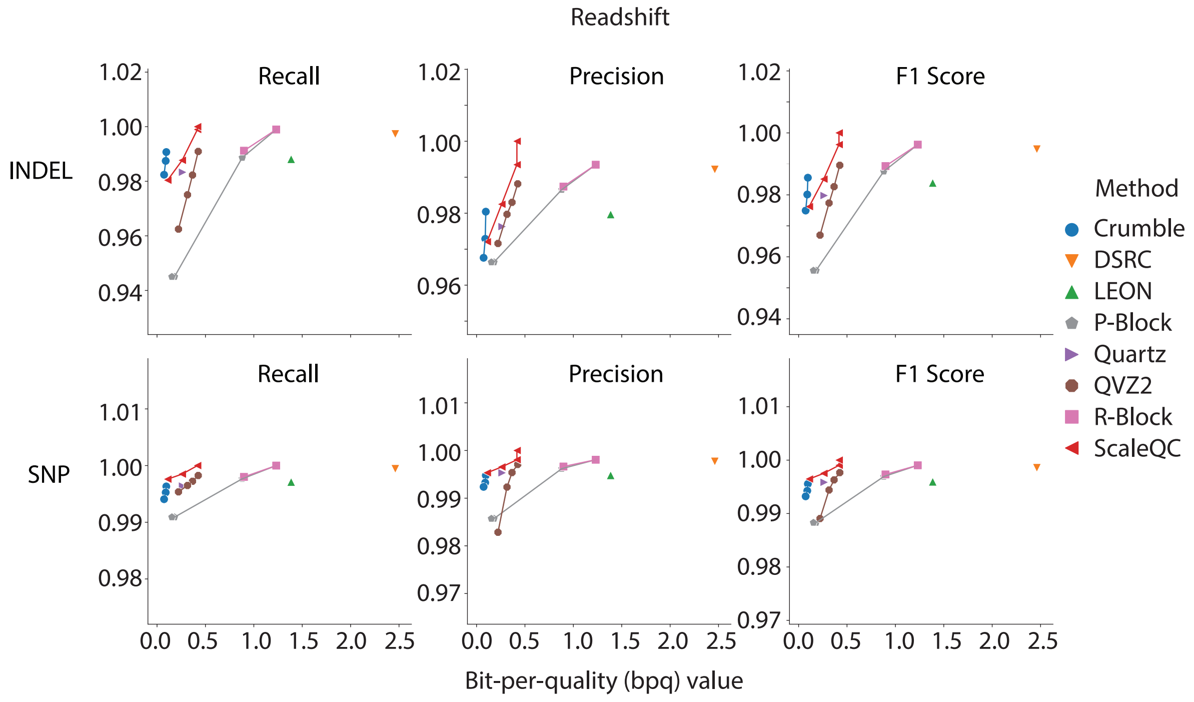


(b) Readshift

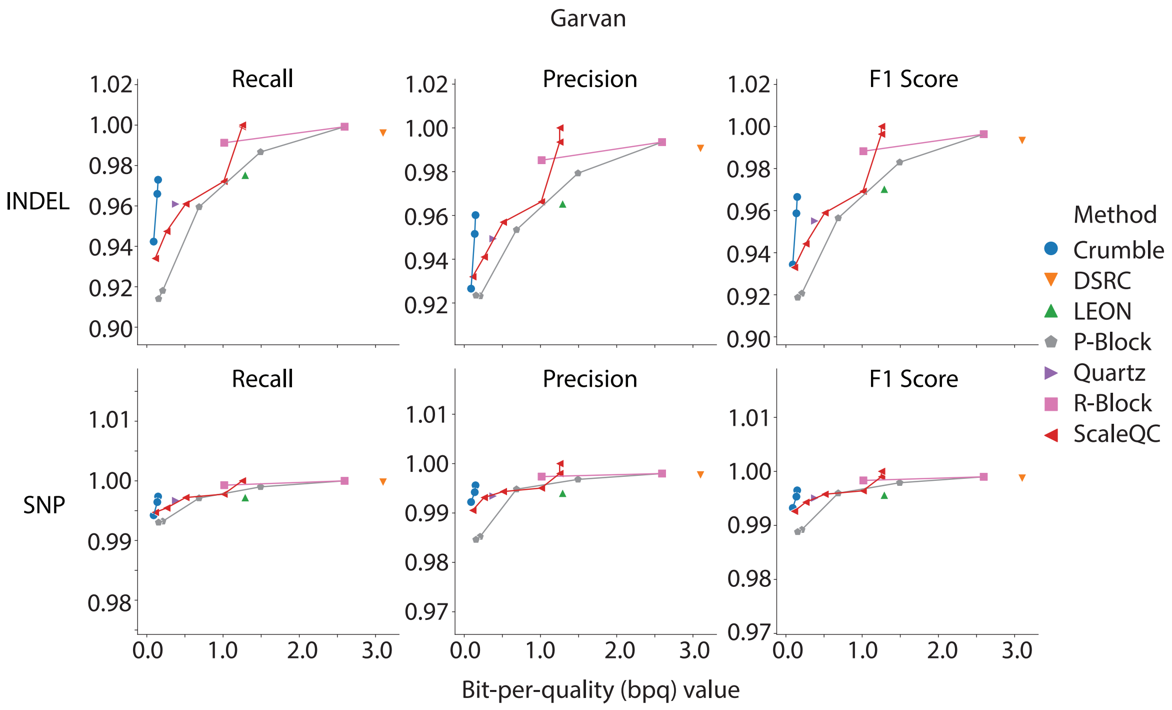


(c) Garvan

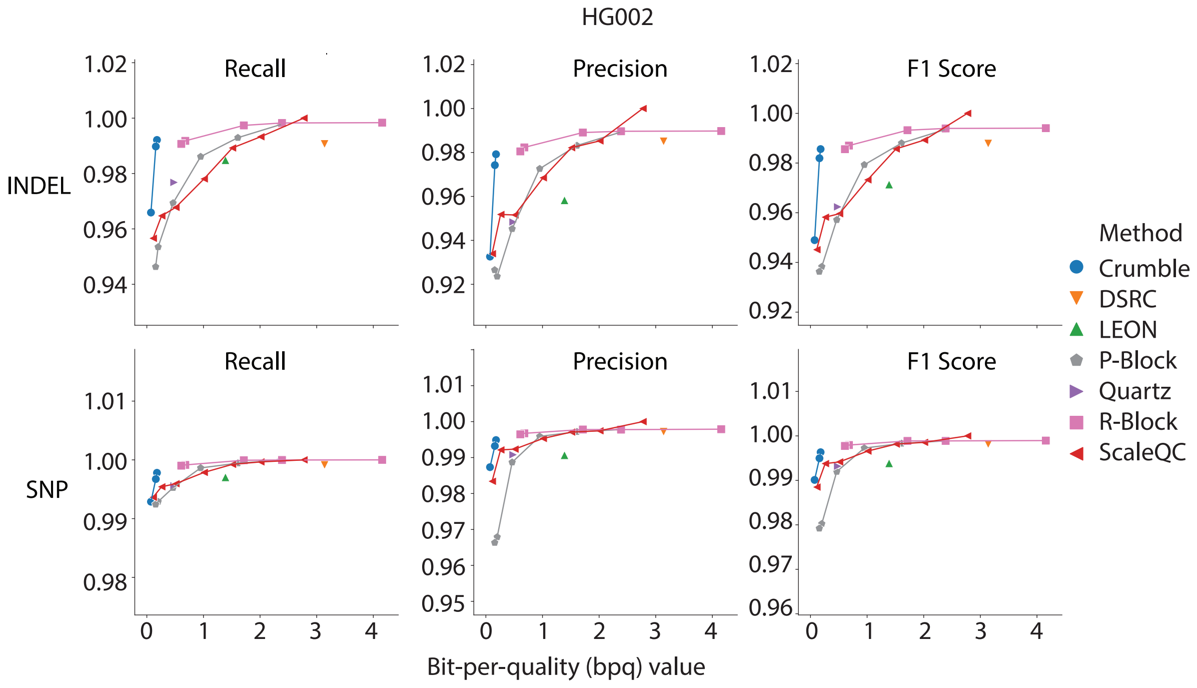


(d) HG002

Figure 1: Comparison of variant calling performance in terms of Recall, Precision and F1 score on decompressed sequencing data for different lossy compressors at different compression ratios. (a) ERR174324, (b) Readshift, (c) Garvan, and (d) HG002.
